# Multidrug-resistant Gram-negative bacteria from patients, hospital environment and healthcare workers: a six-month cross-sectional study

**DOI:** 10.1101/425330

**Authors:** Aline F. R. Sereia, Patricia A. da Cunha, Daniela C. Tartari, Caetana P. Zamparette, Diana A. Estigarribia, Taise C. R. Klein, Ivete Ioshiko Masukawa, Clarice I. Silva, Maria Luiza V. Vieira, Mara C. Scheffer, Dellyana R. Boberg, Ana Paula Christoff, Luiz Felipe V. de Oliveira, Edmundo C. Grisard, Thaís C. M. Sincero

## Abstract

Healthcare-associated infections (HAI) are an important public health threat with the multidrug-resistant (MDR) gram-negative bacteria (GNB) being of particular concern. Here we present the antimicrobial resistance profile of HAI-related GNB (HAIrB) isolated from patients (PT), healthcare workers (HCW) and hospital environment (HE) in a six-month screening program. From the 180 sampling points distributed in six hospital units, a total of 1,080 swabs were collected allowing the isolation of 390 HAIrB: 50.5% from HE, 42.6% from PT and 6.9% from HCW. Among the HAIrB, 32.6% were characterized as MDR and 38.7% as extended-spectrum cephalosporins resistant (ESC-R), showing no differences in the distribution between PT, HE and HCW. Carbapenem resistance (CARB-R) was detected for 17.7% of all HAIrB, being higher among *Acinetobacter* spp. isolates (36.5%), followed by Enterobacteriaceae (14.5%) and *Pseudomonas* spp. (11.8%). Except for the ICU, that revealed higher MDR, CARB-R and ESC-R rates, HAIrB-resistant profiles were similarly detected within the hospital units. Prevalence of *bla*_KPC-like_ and *bla*_CTX-M-1_ β-lactamases-resistance genes was higher in *K. pneumoniae* and *E. cloacae* complex, while *bla*_OXA-23-like_ and *bla*_SPM-like_ were higher in *A. baumannii* and *P. aeruginosa*, respectively. This study reveals that the spreading of HAIrB within a hospital environment is higher than predicted, indicating that healthcare workers, hospital areas and equipment are key players on dissemination of MDR gram-negative bacteria and shows that an active surveillance program can provide precise understanding and direct actions towards control of HAI.

## Introduction

Healthcare-associated infections (HAI) are important public health threats requiring continuous monitoring and efficient surveillance programs (1). HAI caused by multidrug-resistant (MDR) gram-negative bacteria (GNB) are of particular concern, with high-risk global alerts (2-4). HAI can seriously affect patient health, promoting long-term hospital stays and increasing the mortality, in addition to impose high costs for the healthcare system (5-7). There are many evidences that the hospital environment and the healthcare workers are key players on large-scale dissemination of MDR bacteria (8-11). Also, the combination of fast human mobility around the world with selective pressure by overuse and misuse of antibiotics in human and food-producing animals along with the difficulties in adopting simple control measures, form the perfect system to ensure the spread of MDR bacteria (12-14). In this scenario, adoption of surveillance programs based on new technologies associated with the rational management of antimicrobials and the continuous training for healthcare workers can allow an effective control of HAI transmission, ensuring the patient safety and a consequent reduction of direct and indirect healthcare costs (5, 15-16).

To better understand the antimicrobial resistance profile and the dissemination of HAI-related GNB in the healthcare setting we carried out a six-month surveillance program targeting patients, hospital environment and healthcare workers.

## Material and Methods

### Study design

The Healthcare-associated Infections Microbiome Project (HAIMP) was carried out at the Professor Polydoro Ernani de São Thiago University Hospital of Federal University of Santa Catarina (UFSC, Florianópolis/SC, Brazil). The UFSC Human Research Ethics Committee approved this project (number 32930514.0.0000.0121). Between April and September of 2015, a total of 1,080 samples were collected from patients (PT: rectal, nasal and hands swabs; n=198), hospital environment and equipment (HE: high-touch surfaces; n=666) and healthcare workers (HCW: hands, cell phone and protective clothing; n=216). These samples were collected monthly from 180 points (Table S1) distributed in six hospital units: emergency ward (EMG), internal medicine ward (IMW), surgical ward (SUW), general surgery unit (GSU) and intensive care unit A and B (ICU-A and ICU-B). All participants were initially informed about the study aims and sampling was carried out upon a signature of an informed consent. Only long-term hospitalized patients were included in the present study. The samples were collected using Amies agar gel-containing swabs (Copan Inc., Italy) and stored at 4 °C until processing (within 48 hours).

### Phenotypic analyses

Collected swabs were inoculated in Brain Heart Infusion (BHI) broth (BD, USA), incubated for 12-18 hours at 36 °C (±1 °C) prior to plating on selective MacConkey agar (BD, USA), following incubation (18-24 hours) at 36 °C (±1 °C). Different morphotypes of Colony-Forming Units (CFU) were isolated and transferred onto new MacConkey agar plates. Identification and antimicrobial susceptibility test (AST) of each isolate were performed using Vitek^®^2 (BioMérieux Inc., USA) GN ID and AST-N239 cards according to the manufacturer’s instructions. Based on the AST results, the isolates were then classified as “not multidrug-resistant” (Not MDR) or “multidrug-resistant” (MDR), according to the acquired resistance classification (17).

### Genotypic analyses

DNA from all GNB isolates was obtained using a magnetic beads protocol (Neoprospecta Microbiome Technologies, Brazil) and quantified using Qubit dsDNA BR Assay Kit (Invitrogen, USA). A panel of the most important β-lactamases genes in the Brazilian scenario (Table S2) was tested by qPCR using specific primers and hydrolysis probes in a duplex or triplex configuration. The qPCR reactions were carried out in a 10 μL final volume, containing 0.5 ng of DNA and 1X Master Mix (Cy5^®^, HEX^TM^ and FAM^TM^ labeled probes; ROX^TM^ as passive reference; specific forward and reverse primers) (Neoprospecta Microbiome Technologies, Brazil). A negative reaction control and a positive control of each resistance gene (characterized strains containing the gene of interest) were included and the assays were carried out in triplicate. The qPCR amplifications were performed on an ABI 7500 Fast Real-Time PCR System (Applied Biosystems, Foster City, CA, USA), using the following thermal conditions: 95 °C for 10 min, 35 cycles of 95 °C for 15 s, 60 °C for 30 s and 72 °C for 30 s.

Some GNB samples were identified via high-throughput sequencing of 16S rRNA V3/V4 region (Neoprospecta Microbiome Technologies, Brazil) for species confirmation purposes. The 16S rRNA libraries were sequenced using the MiSeq Sequencing System (Illumina Inc. San Diego, CA, USA) with the V2 kit, 300 Cycles, single-end sequencing.

Qui-square or Fisher’s exact test were used for pairwise or proportions comparisons, with significance level set at P < 0.05. All statistical analyses were performed using MedCalc Statistical Software version 14.8.1 (Ostend, Belgium).

## Results

### Gram-negative bacteria (GNB) identification

Among the 1,080 samples collected during the six-month hospital screening, 25.5% (275) were positive for the presence of HAI-related GNB, with statically significant differences between the samples sources (p<0.0001): 53.1% (146) were from hospital environment (HE), 37.5% (103) were from patients (PT) and 9.5% (26) were from healthcare workers (HCW). The highest incidence of positive samples for HAI-related GNB was found in the EMG (32.9%; 77/234), followed by SUW (30.1%; 47/156), ICU-B (25.8%; 51/198), IMW (24.1%; 39/162), ICU-A (23.7%; 44/186) and GSU (11.8%; 17/144); only GSU was statistically different from the other units (p<0.05).

We obtained 390 isolates from the HAI-related GNB positive samples: 197 were from HE (50.5%), 166 from PT (42.6%) and 27 from HCW (6.9%). The phenotypic identification by Vitek^®^2 (GN ID card) allowed the identification of 40 distinct species from 380 isolates (Figure 1, Table S3). Among these, *Escherichia coli, Klebsiella pneumoniae, Pantoea* spp., *Acinetobacter baumannii* complex and *Enterobacter cloacae* complex species were the most abundant, being found at similar frequencies. Identification by Vitek^®^2 GN ID card failed for 10 (2.6%) of the isolates.

**Figure 1.**
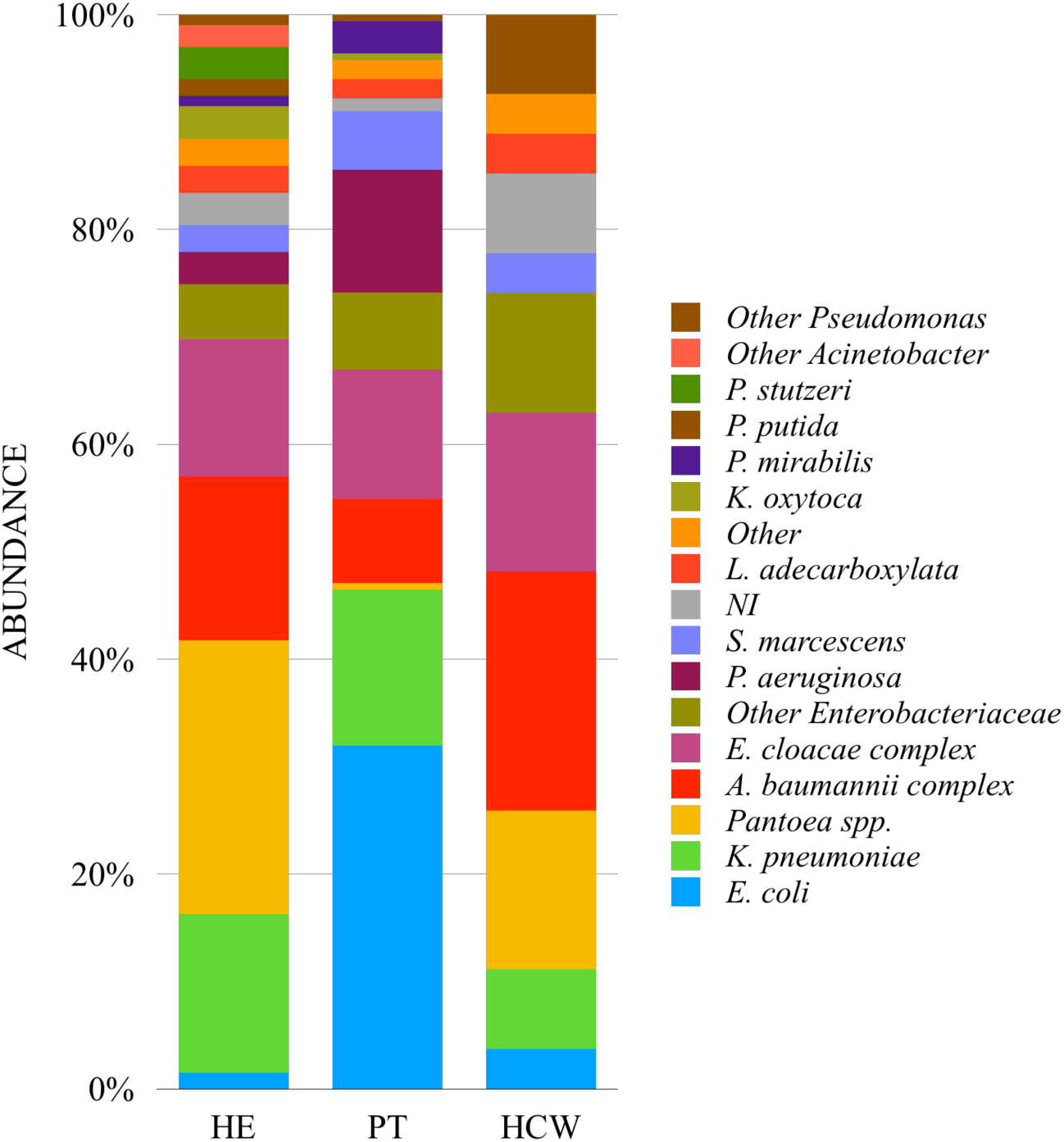
Relative abundance of gram-negative bacteria (GNB) related to healthcare associated infection (HAI) isolated from patients (PT), hospital environment and equipment (HE) and healthcare workers (HCW). NI: Not Identified.

### Antimicrobial Susceptibility Test (AST)

Out of the 390 HAI-related GNB, 310 isolates could be analyzed (Vitek^®^2 AST card failed for 70 and GN ID card failed for 10 of the isolates). The AST analyses allowed us to classify 32.6% (101/310) of the HAI-related GNB as multidrug-resistant (MDR), with no statistically significant difference between the three sources: PT 42.0% (66/157), HE 23.9% (32/134) and HCW 15.8% (3/19). Resistance of GNB to carbapenems (meropenem, ertapenem or imipenem) was identified in 17.7% of the isolates, being significantly higher in PT (24.2%, p=0.036), followed by HCW (15.8%) and HE (10.4%). Resistance of GNB to extended-spectrum cephalosporins (cefepime, ceftazidime or ceftriaxone) was 38.7%, with no statistically significant differences between the sources: 35.0% for PT, 41.0% for HE and 52.6% for HCW (Figure 2). The distribution of MDR HAI-related GNB by each hospital unit, revealed that ICU-B and SUW had the highest proportion, 57.1% (32/56) and 36.5% (19/52), respectively, followed by ICU-A with 30.9% (17/55), IMW with 29.3% (17/58) and EMG with 21.9% (16/73) (in GSU the MDR profile was not identified) (p=0.0001). The ICU-B also showed the highest prevalence of HAI-related GNB resistant to carbapenems, 44.6% (statistically different from the other units), followed by IMW with 20.7%, ICU-A with 16.4%, SUW with 7.7% and EMG with 6.8% (p<0.0001). Resistance to carbapenems was not detected in GNB isolated from GSU. Once again, resistance to extended-spectrum cephalosporins was higher in ICU-B with 60.7% and in IMW with 44.8%, followed by ICU-A with 36.4%, EMG with 31.5%, SUW with 26.9% and GSU with 18.8% (p=0.0014) (Figure 3, Table S4).

**Figure 2.**
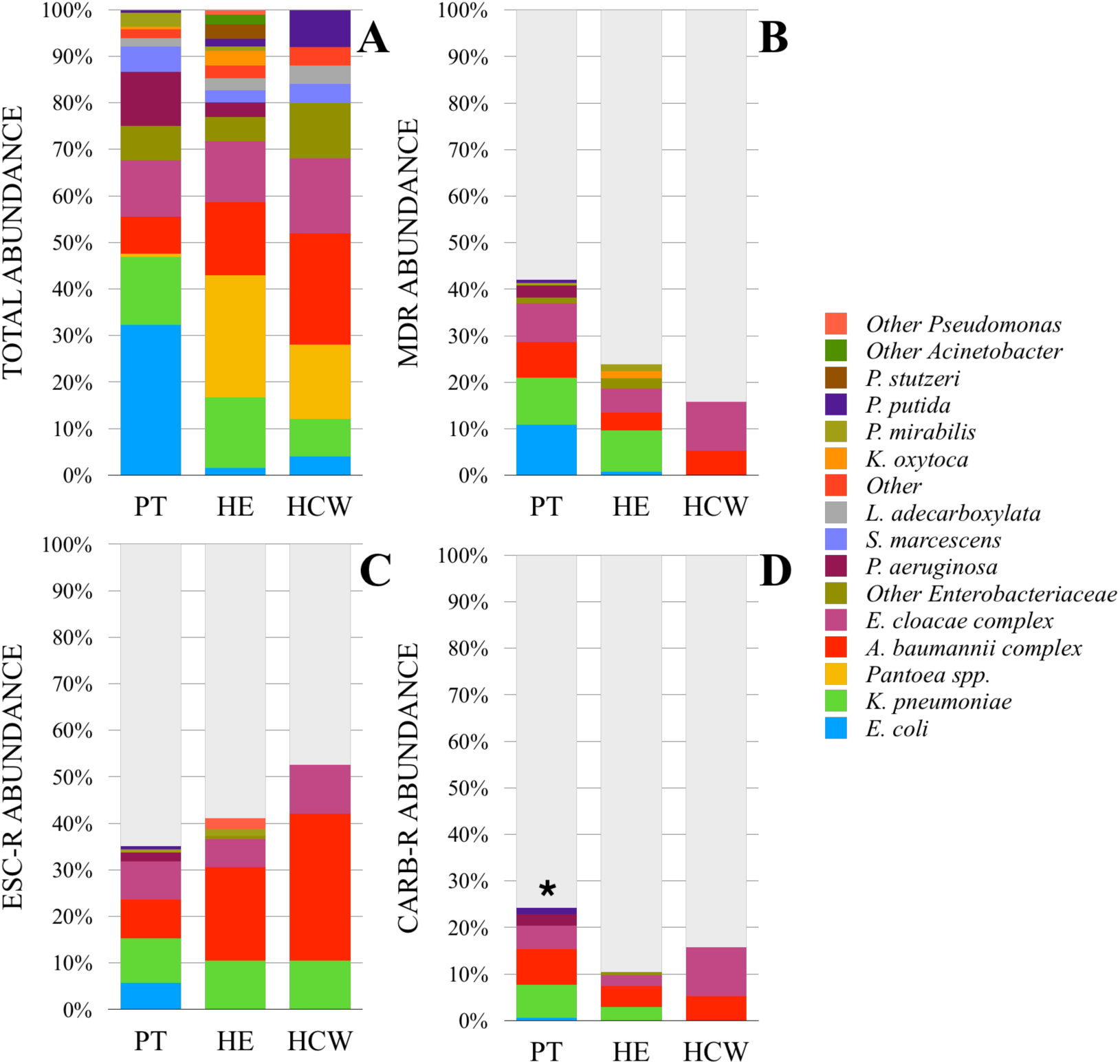
Abundance of Gram negative bacteria according to resistant profile in patients (PT), hospital environment/equipment (HE) and healthcare workers (HCW) (N = 310). A: Total abundance. B: Abundance of multidrug-resistant (MDR) bacteria. C: Abundance of extended-spectrum cephalosporins resistant (ESC-R) bacteria. D: Abundance of carbapenem resistant (CARB-R) bacteria. * p=0.036.

**Figure 3.**
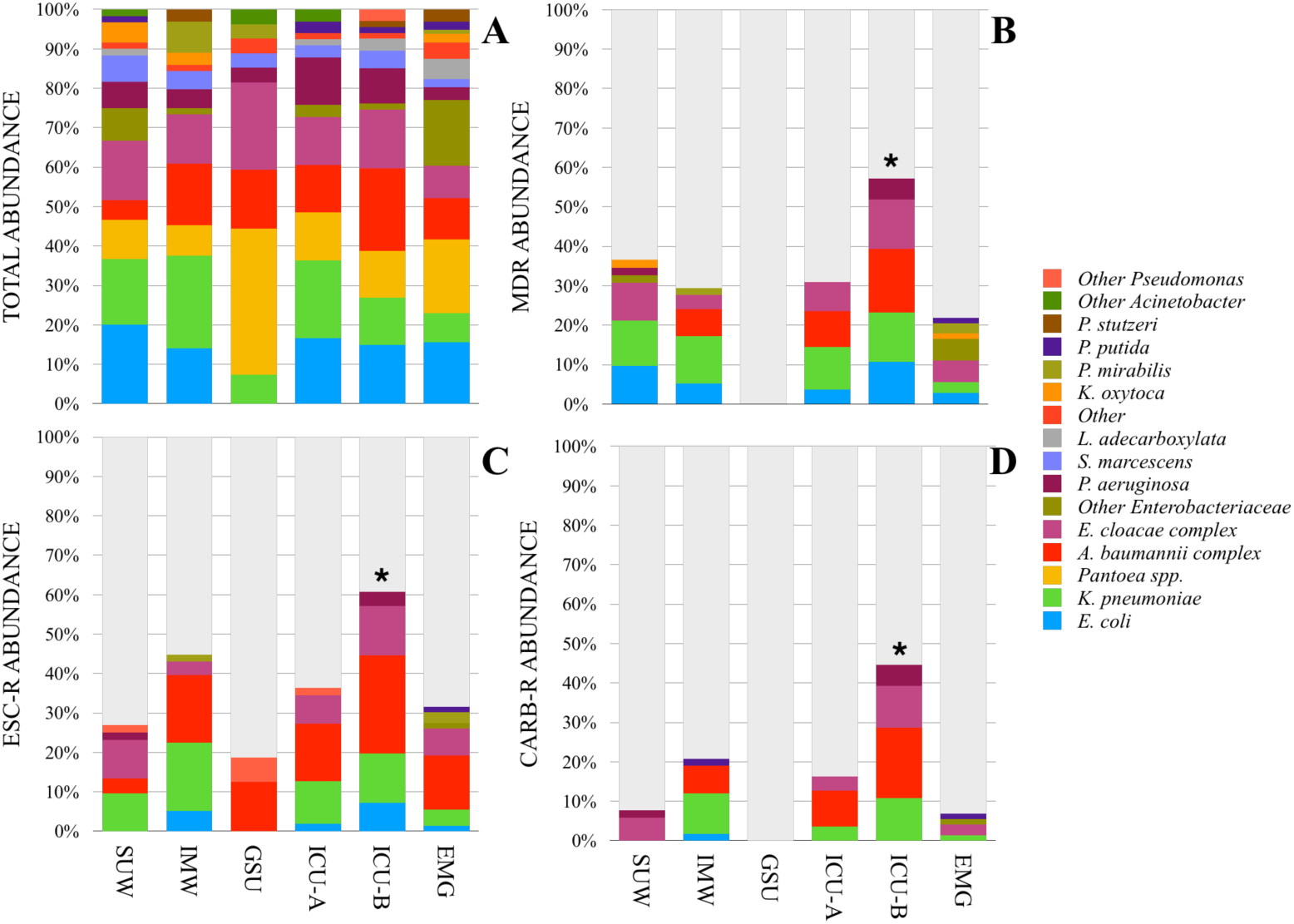
Abundance of Gram negative bacteria according to resistant profile in the six hospital units - Surgical Ward (SUW), Internal Medicine Ward (IMW), General Surgery Unit (GSU), Intensive Care Unit A and B (ICU-A and ICU-B) and Emergency (EMG) (N = 310). A: Total abundance. B: Abundance of multidrug-resistant (MDR) bacteria. C: Abundance of extended-spectrum cephalosporins resistant (ESC-R) bacteria. D: Abundance of carbapenem resistant (CARB-R) bacteria. * p<0.05.

We separated the collected samples into groups (representing the areas of the hospital or the isolation sites) with the distribution of GNB and their resistance profile (Figure 4, Table S5). Analysis of this data allowed us to identify the critical points of contamination within the studied areas.

**Figure 4.**
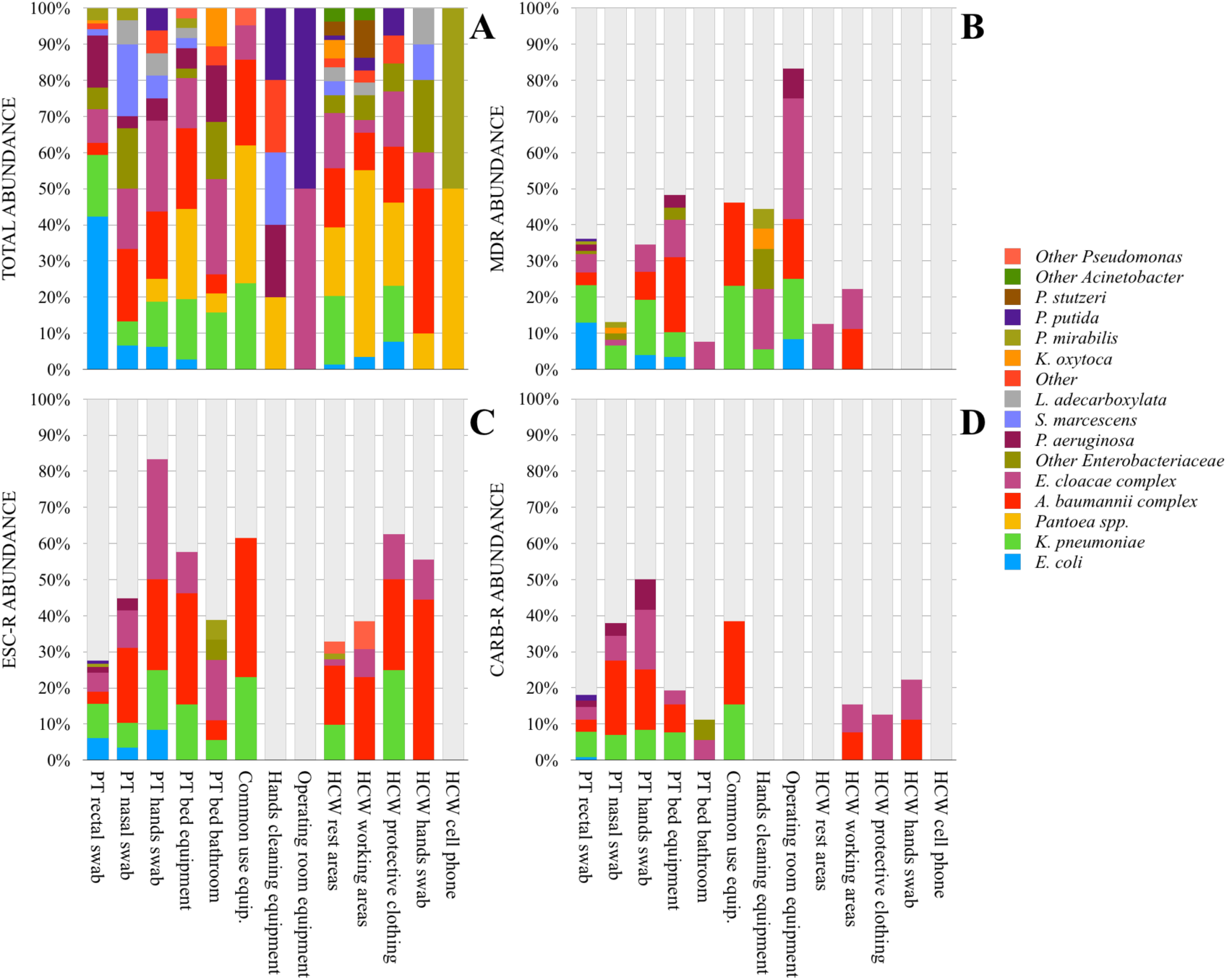
Abundance of Gram negative bacteria according to resistant profile in the different areas of the hospital or the isolation sites (N = 310). A: Total abundance. B: Abundance of multidrug-resistant (MDR) bacteria. C: Abundance of extended-spectrum cephalosporins resistant (ESC-R) bacteria. D: Abundance of carbapenem resistant (CARB-R) bacteria.

### Characterization of the patient-related samples

During the sampling period, a total of 66 patients had their samples collected, including samples from their respective beds (HE samples) in all hospital units, except GSU. The average age of the patients was 63.2 years, being 47.0% female and 53.0% male. The average length of hospital stay was 15.3 days and, unexpectedly, the longer hospitalization was seen for SUW patients, with 24.3 days, followed by IMW (23.4 days), ICU-B (15.0 days), ICU-A (10.4 days) and EMG (6.8 days). More than half of these patients (54.5%) were admitted to more than one unit of the hospital and 65.2% undergone at least one invasive procedure. Medical records also show that 50.0% of the patients did not present HAI during the current hospitalization, 43.9% had a HAI during the current hospitalization and 6.1% were colonized by HAI-related bacteria. The results of the present study showed that 62 patients (93.9%) had at least one HAI-related GNB isolated from their samples and 63.6% of the patients had at least one MDR GNB, 39.4% had an extended-spectrum cephalosporin GNB and 28.8% had a carbapenem resistant GNB. From the patients with two or more sites colonized by HAI-related GNB (28/66 - 46.7%) we identified MDR GNB of same species in 42.9% of the cases, with 35.7% showing an AST very similar (equal MDR and carbapenem resistance classification). Correspondence of HAI-related GNB isolated from patients and from their beds was positive for 31 cases: 16.1% revealed to be the same GNB species with equal MDR and carbanepem resistance profiles.

### Enterobacteriaceae antimicrobial resistance profile

The most common Enterobacteriaceae (n=279) found were *Escherichia coli* (20.4%), *Klebsiella pneumoniae* (19.7%) *Pantoea* spp. (19.4%), *Enterobacter cloacae* complex (17.5%) and *Serratia marcescens* (5.4%).

A total of 220 of the 279 Enterobacteriaceae presented results for the AST. AST results for the three most frequent species of HAI-related Enterobacteriaceae (*Klebsiella pneumoniae, Enterobacter cloaceae* and *Escherichia coli)* in PT, HE and HCW are presented in Tables S6, S7 and S8. Analysis of the antimicrobial resistance profiles did not show statistically significant differences among the different GNB sources (PT, HE and HCW) (Figure 5A). MDR profile was found in 35.0% of the Enterobacteriaceae, carbapenem resistance was present in 14.5% and extended-spectrum cephalosporins resistance was found in 30.9%. In PT, HE and HCW, MDR was found in 40.3% (50/124), 29.4% (25/85) and 18.2% (2/11) of the Enterobacteriaceae isolates, carbapenem resistance was identified in 17.7%, 9.4% and 18.2% and extended-spectrum cephalosporins resistance was present in 31.4%, 29.4% and 36.4%, respectively, with no statistically significant differences among the groups tested. When the three most frequent species of Enterobacteriaceae were analyzed separately, in patients (PT) we found that MDR, carbapenem resistance and extended-spectrum cephalosporins resistance for these species were, respectively, 66.7%, 45.8% and 62.5% for *Klebsiella pneumoniae* (n=24), 65.0%, 40.0% and 65.0% for *Enterobacter cloacae* complex (n=20) and 32.1%, 1.9% and 17.0% for *Escherichia coli* (n=53). For hospital environment (HE) samples, *K. pneumoniae* (n=29) and *E. cloacae* complex (n=25) were classified as MDR in 41.4% and 28.0% of the cases, respectively, 13.8% and 12.0% were carbapenem resistant and 48.3% and 32.0% presented resistance to extended-spectrum cephalosporins. We found a case of a IMW patient colonized by MDR (carbapenem and extended-spectrum cephalosporins resistant) *K. pneumoniae* in the three isolations sites (rectal swab, hands and nasal). In healthcare workers (HCW) we found two *K. pneumoniae*, three *E. cloacae* complex and one *E. coli*, being the two *E. cloacae* complex isolates MDR and resistant to carbapenems.

**Figure 5.**
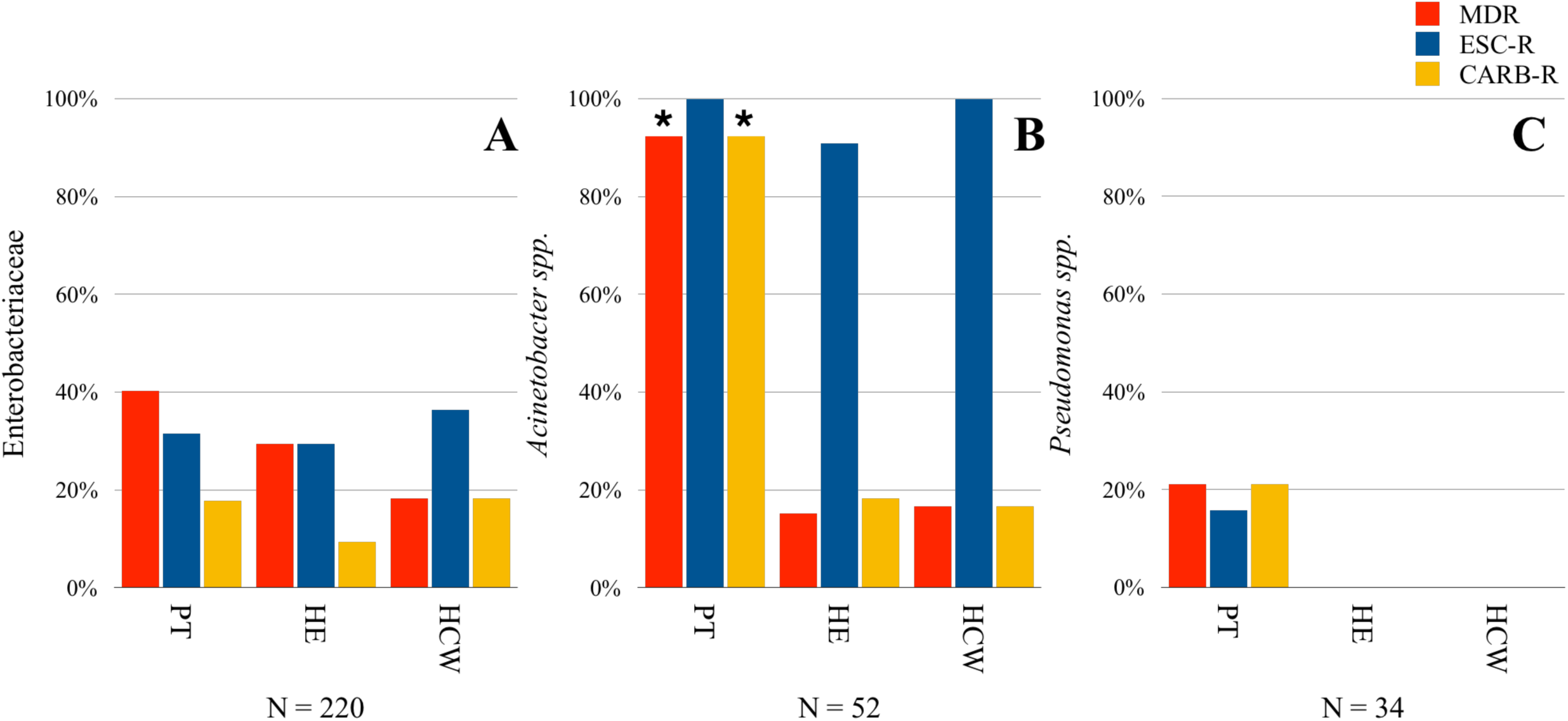
Proportion of multidrug-resistant (MDR), extended-spectrum cephalosporins resistant (ESC-R) and carbapenem resistant (CARB-R) bacteria in patients (PT), hospital environment/equipment (HE) and healthcare workers (HCW). A: Proportions for Enterobacteriaceae; B: Proportions for *Acinetobacter* spp.; C: Proportions for *Pseudomonas* spp. *p<0,0001.

Frequencies of resistance genes detected by qPCR in HAI-related Enterobacteriaceae from PT, HE and HCW are presented in Figure 6 as a heat map, emphasizing the most important GNB of the family (*K. pneumoniae, E. cloacae* complex and *E. coli)* (Table S9 shows resistance genes frequencies with the respective Cq mean). The *bla*_KPC-like_ gene was found in 5.9% (13/220) of Enterobacteriaceae, all of them resistant to carbapenems and identified as *K. pneumoniae, E. cloacae* complex or *Escherichia coli*. In another Enterobacteriaceae species resistant to carbapenems we did not found *bla*_KPC-like_ or any other tested carbapenemase encoding gene. *Klebsiella pneumoniae* recovered from PT (n=24) presented concern beta-lactamase genes frequencies: 91.7% for *bla*_SHV-like_, 37.5% for *bla*_CTX-M-1_ group, 33.3% for *bla*_KPC-like_, 25.0% for *bla*_CTX-M-9_ group and 16.7% for *bla*_CTX-M-8_ group (some of these *K. pneumoniae* harboring three or more beta-lactamase genes).

**Figure 6.**
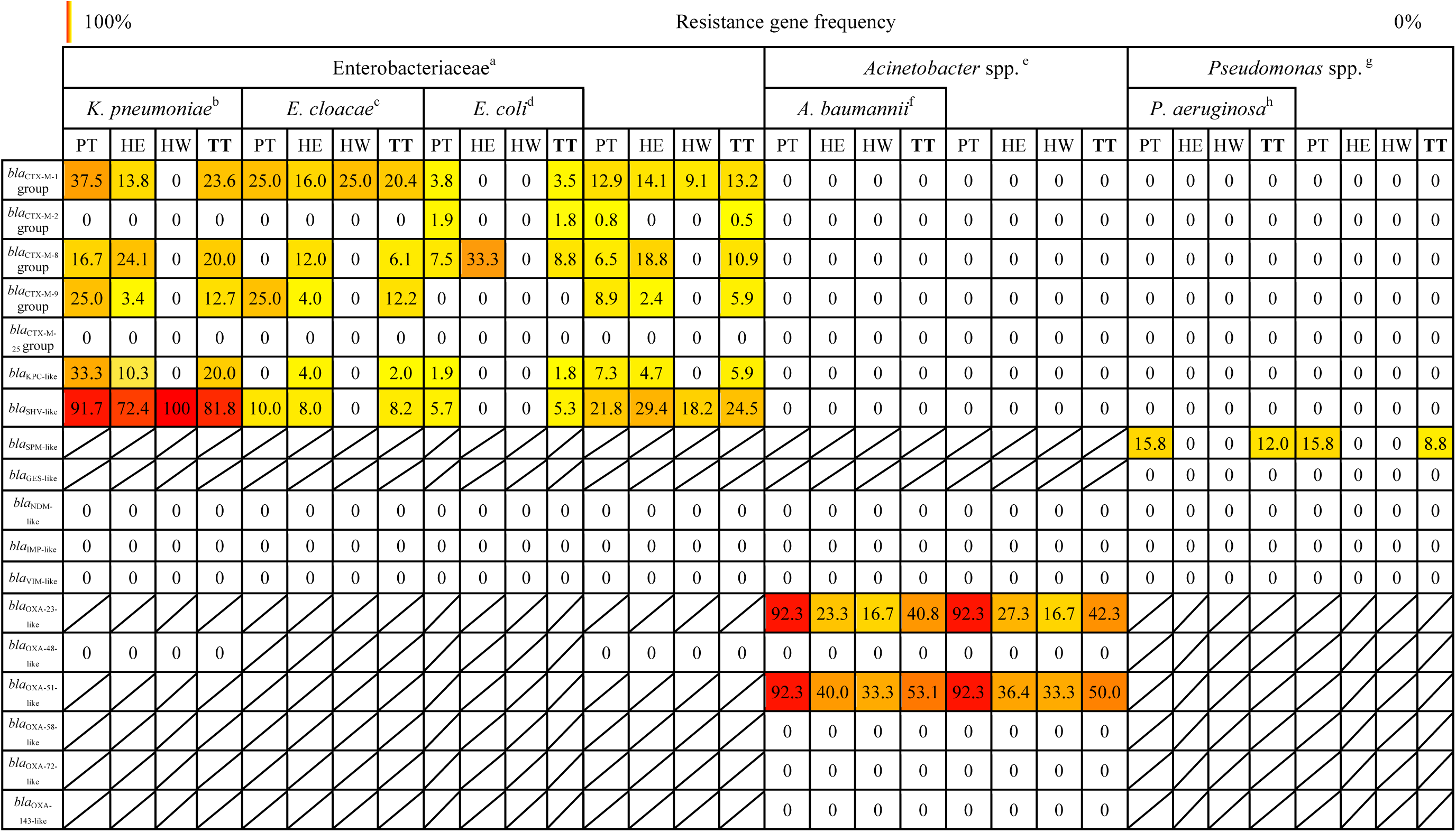
Heat map showing the frequencies of the beta-lactamases resistance genes identified in the total (TT) samples of each gram-negative bacteria (GNB) related to healthcare associated infection (HAI) group and for the most important GNB isolated from patients (PT), hospital environment and equipment (HE) and healthcare workers (HW). The red color indicates high beta-lactamases genes frequencies, while yellow and white colors indicates low and null frequencies, respectively. Cells with cross line represents untested genes. For comparison purposes, we considered only the GNB with Antimicrobial Susceptibility Test (AST). ^a^Enterobacteriaceae with AST = 220. *^b^Klebsiella pneumoniae* with AST = 55. ^c^*Enterobacter cloacae* complex with AST = 49. *^d^Escherichia coli* with AST = 57. ^6^*^e^cinetobacter* spp. with AST = 52. *^f^Acinetobacter baumannii* complex with AST = 49. *^h^Pseudomonas* spp. with AST = 34. ^h^*Pseudomonas aeruginosa* with AST = 25.

### *Acinetobacter* spp. antimicrobial resistance profile

Using the phenotypic identification, we found that 92.5% (49/53) were *Acinetobacter baumannii* complex, 5.7% (3/53) were *Acinetobacter lwoffii* and 1.9% (1/53) *Acinetobacter radioresistens*.

Table S10 shows the phenotypic antimicrobial resistance profile of 52 HAI-related *Acinetobacter* spp. in PT, HE and HCW (Table S11 shows AST results for *Acinetobacter baumannii* complex separately). For *Acinetobacter* spp. the highest resistance to antimicrobials was seen in PT (Figure 5B): 92.3% (12/13) were MDR and resistant to carpapenems (imipenem or meropenem). The *bla*_OXA-23-like_ gene, which confers resistance to carbapenems, was identified at the same frequency (92.3%) (Figure 6). In HE and HCW carbapenem resistance were lower, 18.2% (6/33) and 16.7% (1/6) respectively. When PT, HE and HCW were analyzed together *Acinetobacter* spp. still showed the highest carbapenem resistance of all HAI-related GNB: 36.5%. MDR profile was found in 34.6% of *Acinetobacter* spp.: 92.3% (12/13) in PT, 15.2% (5/33) in HE and 16.7% (1/6) in HCW. The carbapenems resistance and MDR profiles were statistically significant different when we compare PT with HE or HCW (p<0.0001). Extended-spectrum cephalosporins resistance (cefepine, ceftazidime or ceftriaxone) was found in 100%, 90.9% and 100% of PT, HE and HCW respectively (Figure 5B).

Figure 6 shows the frequencies of beta-lactamase genes detected by qPCR in HAI-related *Acinetobacter* spp. from PT, HE and HCW, with *A. baumannii* complex analyzed separately (Table S9). In *A. baumannii* complex (n=49) the *bla*_OXA-23-like_ was found in 40.8% of the isolates and, interestingly, *bla*_OXA-51-like_ (intrinsic in *A. baumannii)* was found only in 53.1. Because of this unexpected observation, all *Acinetobacter* spp. were identified by high-throughput sequencing of V3/V4 region of the 16S rRNA for species confirmation. The isolates identified *as A. baumannii* complex by Vitek^®^2, GN ID card, with no detection of *bla*_OXA-51-like_ gene (n=23), were identified by 16S rRNA sequencing as *A. baumannii* (one), *A. calcoaceticus* (seven) and *A. pittii, A. soli* or *A. nosocomialis* (15), which explains the absence of the gene in most of the cases.

### *Pseudomonas* spp. antimicrobial resistance profile

For *Pseudomonas* spp. isolated in the survey (n=39), 64.1% were identified as *Pseudomonas aeruginosa*, 15.4% were *Pseudomonas putida*, 15.4% were *Pseudomonas stutzeri* and 5.2% were *Pseudomonas luteola* or *Pseudomonas oryzihabitans*.

AST profile of HAI-related *Pseudomonas* spp. from PT and HE can be verified in Table S12 (Table S13 shows AST results for *Pseudomonas aeruginosa* separately). This GNB group presented the more susceptible antimicrobial profiles, with resistance to important classes, like carbapenems and antipseudomonal cephalosporins (cefepine or ceftazidime), identified only in PT *Pseudomonas* spp. (n=19): 21.1% and 15.8%, respectively. The acquired resistance classification showed that 15.8% of *Pseudomonas* spp. in PT were MDR (Figure 5C) and presented the *bla*_SPM-like_ gene, that classically confers resistance do carbapenems (Figure 6, Table S9). All *Pseudomonas* spp. were 100% susceptible to Colistin.

Antimicrobial resistant profiles (MDR or carbapenem resistance) and *bla*_SPM-like_ were found only in *Pseudomonas aeruginosa* and is important to point out that three of them were isolated from the same ICU patient (one *P. aeruginosa* for each isolation site - hand, nasal and rectal swabs).

## Discussion

There is a high risk of acquiring HAI in developing countries, reaching up to 27%. The fight to improve patient safety is difficult because of the increasing antimicrobial resistance rates, coupled to other serious health systems problems and to the fact that health authorities are not sufficiently prepared to face the problem (18-19). Gram-negative bacteria (GNB) present a good performance against the antibiotics, making them a leading cause of HAI and a matter of great concern for the currently available therapies (20).

Here, we presented the first results for culture-dependent samples of the Healthcare-associated Infections Microbiome Project (HAIMP) that has been carried out in a teaching hospital in Southern Brazil. During six months, patients (PT), healthcare workers (HCW) and hospital environment (HE) were monitored using swab samples for the screening of HAI-related gram-negative bacteria (GNB).

A total of 390 GNB were isolated during the surveillance program. Seven species/genus (or species complex) accounted for 80% (304/380) of the total GNB identified, almost all of them classically identified in previous healthcare studies, like *Klebsiella penumoniae, Enterobacter cloacae* complex and *Acinetobacter baumannii* complex (21).

From all the collected swabs, the samples from PT, HE and HCW had different frequencies of GNB. There is a higher contamination rate of the patients and hospital environment samples by HAI-related GNB, when compared to the healthcare workers. Giving the inclusion criteria applied in our study, more than 90% of the patients presented a HAI-related GNB. We found several cases where the same GNB species (with similar or equal resistance profile) was identified in samples from the patients and their room, suggesting a cross contamination and demonstrating the importance of the hospital environment in the HAI dissemination (11, 22-25). Patients rectal swabs were the site with the highest positivity, followed by nasal and hands swabs. However, GNB from nasal and hands swabs had relatively higher resistance rates than GNB from rectal swabs, probably due to different composition of bacterial populations (26), making them more likely to be colonized by MDR GNB. Nasal colonization studies usually emphasize gram-positive bacteria (27, 28), however it is important to highlight the high contamination rate by GNB identified in nasal swab samples of this survey: 23 of 66 participating patients had at least one GNB isolated from this collection site, some of them MDR and resistant to carbapenems.

We found that 32.6% of the HAI-related GNB were MDR and no statistically significant differences were seen for PT (47.7%), HE (29.1%) and HCW (26.3%). Brazilian studies found MDR profile for GNB isolated from patients ranging from 32% to 48% (29, 30).

In the present study, the resistance rate to carbapenems was 17.7%, being 24.2% when considering only the PT samples. The average carbapenem resistance reported in the literature for HAI diagnosed patients samples is 42.7% (31-37). The extended-spectrum cephalosporins resitance were identified in 38.7% of the total GNB (35.0% for PT). Additionally, the average extended-spectrum cephalosporins resistance or ESBL profile reported from the literature for patient-related samples was 31.7% (21, 32, 38-40).

The results presented for hospital environment and equipment showed that the rest areas of the healthcare workers, like the lunch and the sleeping rooms, were highly contaminated, also including positive results for MDR bacteria. The number of isolated GNB found in these areas were only smaller than those from the rectal swabs. Common work areas and hospital medical equipment were also critical points of contamination, many many harbouring carbapenems-resistant GNB. We identified five cases of healthcare workers contamination with MDR GNB (four from hands e one from protective clothes), three of them resistant to carbapenems.

The only hospital unit that showed statistically different frequencies of HAI-related GNB was GSU, with a low frequency of GNB. This result could be due to the fact that no patient samples were collected in this unit and also because one of the surgical rooms was always collected after its disinfection. Among the other hospital units we saw similar GNB rates, indicating a systematic contamination in the hospital. Additionally, PT, HE and HCW isolates did not show significant differences in the resistance profile, which suggests a homogeneous spread of resistant GNB through the hospital.

ICU-B had the most concerning results for the antimicrobial profile of HAI-related GNB, with the highest frequencies of MDR, carbapenem and extended-spectrum cephalosporins resistance when compared to the others units. Rubio *et al*. (21) had also found a significant difference between MDR GNB isolated from ICU patients and non-ICU patients. A possible explanation for this results is that the ICU-B is an adult ICU that receives critically ill patients, with longer length of stay, which exposes patients, healthcare workers and the environment to an increased chance of contamination by multi-resistant bacteria. After the ICU-B, IMW and ICU-A showed the highest carbapenems and extended-spectrum cephalosporins resistance rates.

Enterobacteriaceae showed very similar resistance profiles among the three tested groups: PT, HE and HCW. The profile identified reinforces the systematically spread of GNB occurring in patients, healthcare workers and environment. Within the most important species of the family, *Escherichia coli* presented a more susceptible antimicrobial profile and *Klebsiella pneumoniae* and *Enterobacter cloacae* complex presented the the highest rates of MDR and carbapenem resistance. The AST profile identified here for *E. coli* was very similar to previous studies (32, 33). The AST for *K. pneumoniae* isolated from environment and healthcare workers was very similar to the AST identified for *Klebsiella* spp. isolated from patients in previous studies (32, 34). However, when we compare the AST results of *K. pneumoniae* isolated from patients in the present study we found higher resistance rates than previous reports (32, 34). For *E. cloacae* complex we found a similar scenario: in the present study the isolates from patients had higher resistance rates than reported in literature (32, 34). It is important to emphasize that the study presented here includes GNB isolated from patients with infection or colonization, as well as from the environment and healthcare workers, which further draws attention to the higher resistance rates that were found.

The *bla*_SHV-like_ shows the highest frequencies of all resistance genes tested, reaching up 91.7% for *K. pneumoniae* isolated from patients, and the frequencies found (in Enterobacteriaceae or specific species) are in accordance with previous studies (41, 42, 43). The *bla*_CTX-M-8_, *bla*_CTX-m-8_ and *bla*_CTX-M-9_ groups were the most frequents of the CTX-M family genes. These findings were similar to several studies (42, 44). The *bla*_CTX-M-2_ was identified with lower frequencies and only in *E. coli*, as previously reported (38, 45). However, other studies reported higher *bla*_CTX-M-2_ frequencies in Enterobacteriaceae, with results ranging between 52.3% and 89.3% (41, 42, 46). For the *bla*_KPC-like_ we found a 6.4% frequency for Enterobacteriaceae and 33.3% for *K. pneumoniae* isolated from patients, in accordance with previous studies (38, 47-50). In *Acinetobacter* spp. we found an antimicrobial profile significantly more resistant in patients than in hospital settings and healthcare workers.

The *bla*_OXA-23-like_ detection in *A. baumannii* complex isolated from patients followed the rate of carbapenem resistance, actually they were the same (92.3%), which shows the importance of this carbapenemase gene for this species. The carbapenem resistance studies with *Acinetobacter* spp. isolated from patients ranged between 30.0% and 91.2% (30, 31, 34-36), while *bla*_OXA-23-like_ frequencies are reported from 41.7% to 100% (51, 52). The absence of *bla*_OXA-51-like_ in some *A. baumannii* complex isolates, that includes *A. baumannii, A. calcoaceticus, A. pittii* and *A. nosocomialis* species (53), allowed us to show the correct species identification of 22 *non-baumannii* isolates by V3/V4 16S rRNA sequencing. A study conducted by Vasconcelos *et al*. (54) reported the absence of *bla*_OXA-51-like_ in *non-baumannii* isolates, while Teixeira *et al*. (55) reported a frequency of 41.7% for *non-baumannii*. We did not find the *bla*_OXA-143-like_ gene and the studies report absence (54), low (56) and high frequencies for this carbapenemase gene (52). Some other studies show null or low prevalences of *bla*_OXA-58-like_ (31, 57) and *bla*_OXA-72-like_ (54, 58), while they were not identified in the present study.

The AST profile for *Pseudomonas aeruginosa* isolated from patients identified here was similar to previous reported by Jones *et al*. (32), while Gales *et al*. (34) found a higher resistance rate for all the tested antibiotics. However, in both studies we see higher carbapenems resistance rates, that ranged from 38.4% to 46.7%. In the present study we identified the resistance to meropenem was found in 21.1% and to imipenem in 26.3% of the *P. aeruginosa* isolated from patients.

In *P. aeruginosa*, we highlight the presence of *bla*_SPM-like_ gene (15.8% for isolates from patients) which highly accounted to define the phenotypic MDR profile. Previous studies with *P. aeruginosa* reported *bla*_SPM-like_ frequencies close to 6.0% (33, 36), while studies with *P. aeruginosa* with a MDR or carbapenem resistant profile reported higher and more variable frequencies, ranging from 17.8% to 64.1% (59, 60).

Despite the report of several cases of *bla*_NDM-like_ in Brazil in different GNB species (31, 61), the present study did not identify this carbapenemase gene, which can be explained by its low incidence in the country, 0.97% in Enterobacteriaceae according to Rozales *et al*. (62). The *bla*_IMP-like_ and *bla*_VIM-like_ metalo-beta-lactamases (MBL) genes and *bla*_GES-like_ carbapenemase gene were also not found in the present surveillance program, however others Brazilian studies found low frequencies of these genes in *P. aeruginosa* and *A. baumannii* (33, 36, 43, 59).

It is important to point out that all studies cited were carried out with GNB isolated from patients and the identification of the resistance genes were restricted in most of the cases to resistant GNB (ESBL, MBL or carbapenem resistant, for example), becoming an important bias for the comparative analysis.

There were two interesting cases of patients showing contaminated by GNB in the three isolation sites (hands, nasal and rectal swabs). One of them was a case involving *P. aeruginosa* that occurred to an ICU patient and the other was a IMW patient, colonized by *K. pneumoniae*. In both cases the GNB was present in the three isolation sites and had the same MDR, carbapenem and extended-spectrum cephalosporins resistant profile. These findings help to demonstrate the importance of hygiene practices and patients surveillance, with special attention to the barrier-breaking events, since the contamination of isolation sites like the hands, facilitates the contamination of bed equipments and healthcare workers with MDR GNB, as well as could difficult the patient treatment, once a recolonization event can occur. For our knowledge, this is the first extensive study for the screening and antimicrobial resistance profile characterization of the HAI-related GNB in a hospital that sought the correlation among patients, environment and healthcare workers. This study helped to understand more about resistant HAI-related GNB circulating in a hospital and reveals that spread of HAI-related GNB within the hospital environment is higher than predicted, indicating that healthcare workers, hospital areas and equipment are key factors on MDR gram-negative bacteria dissemination. An active surveillance program can be a powerful tool for the understanding and control of HAI, however, it is important to point out that the biggest challenge to be faced with the implementation of an active surveillance program is that it requires a formal leadership and a committed team to use well all their potential results.

## Acknowledgments

We thank Dr. Ana Cristina Gales (*ALERTA Laboratory, Federal University of São Paulo)* who kindly provided control strains for the study.

